# Local genetic effects on gene expression across 44 human tissues

**DOI:** 10.1101/074450

**Authors:** François Aguet, Andrew A. Brown, Stephane E. Castel, Joe R. Davis, Pejman Mohammadi, Ayellet V. Segrè, Zachary Zappala, Nathan S. Abell, Laure Frésard, Eric R. Gamazon, Ellen Gelfand, Michael J. Gloudemans, Yuan He, Farhad Hormozdiari, Xiao Li, Xin Li, Boxiang Liu, Diego Garrido-Martín, Halit Ongen, John J. Palowitch, YoSon Park, Christine B. Peterson, Gerald Quon, Stephan Ripke, Andrey A. Shabalin, Tyler C. Shimko, Benjamin J. Strober, Timothy J. Sullivan, Nicole A. Teran, Emily K. Tsang, Hailei Zhang, Yi-Hui Zhou, Alexis Battle, Carlos D. Bustamonte, Nancy J. Cox, Barbara E. Engelhardt, Eleazar Eskin, Gad Getz, Manolis Kellis, Gen Li, Daniel G. MacArthur, Andrew B. Nobel, Chiara Sabbati, Xiaoquan Wen, Fred A. Wright, GTEx Consortium, Tuuli Lappalainen, Kristin G. Ardlie, Emmanouil T. Dermitzakis, Christopher D. Brown, Stephen B. Montgomery

## Abstract

Expression quantitative trait locus (eQTL) mapping provides a powerful means to identify functional variants influencing gene expression and disease pathogenesis. We report the identification of cis-eQTLs from 7,051 post-mortem samples representing 44 tissues and 449 individuals as part of the Genotype-Tissue Expression (GTEx) project. We find a cis-eQTL for 88% of all annotated protein-coding genes, with one-third having multiple independent effects. We identify numerous tissue-specific cis-eQTLs, highlighting the unique functional impact of regulatory variation in diverse tissues. By integrating large-scale functional genomics data and state-of-the-art fine-mapping algorithms, we identify multiple features predictive of tissue-specific and shared regulatory effects. We improve estimates of cis-eQTL sharing and effect sizes using allele specific expression across tissues. Finally, we demonstrate the utility of this large compendium of cis-eQTLs for understanding the tissue-specific etiology of complex traits, including coronary artery disease. The GTEx project provides an exceptional resource that has improved our understanding of gene regulation across tissues and the role of regulatory variation in human genetic diseases.

## Introduction

Genome-wide association studies (GWAS) have identified a wealth of genetic variants associated with complex traits and disease risk. However, characterizing the molecular and cellular mechanisms through which these variants act remains a major challenge that limits our understanding of disease pathogenesis and the development of therapeutic interventions. Expression quantitative trait locus (eQTL) studies provide a systematic approach to characterize the molecular consequences of genetic variation across tissues and cell types^1–4^. Multiple studies have identified eQTLs for thousands of genes^5–7^, providing novel insights into gene regulation and enabling the interpretation of GWAS signals^8–12^. These studies have largely been performed in a few easily accessible cell types and cell lines, precluding interpretation of the systemic and tissue-specific consequences of genetic variation. To overcome these limitations, the Genotype Tissue Expression (GTEx) project was designed to identify and characterize eQTLs across a broad range of tissues. During the pilot phase, which focused on nine tissues, the GTEx project highlighted patterns of eQTL tissue-specificity and demonstrated the value of multi-tissue study designs for identifying causal genes and tissues for trait-associated variants^1^. These results indicated that the identification of eQTLs across an even broader range of tissues would drastically improve characterization of the gene- and tissue-specific consequences of genetic variants.

Here, we report on the discovery of cis-eQTLs across an expanded collection of 44 tissues in the GTEx V6p study. This dataset consists of 7,051 transcriptomes from 449 individuals and 44 tissues (median 16 tissues per individual, 127 samples per tissue), including multiple tissues that are difficult to sample such as 10 distinct brain regions. With this dataset, we identified cis-eQTLs within each tissue and characterized the sharing of eQTLs across tissues. We next assessed the relationship between tissue-specific and shared eQTLs with different functional annotations, including promoters, enhancers and Hi-C contacts, and with allele-specific expression (ASE). Finally, we demonstrated the utility of this multi-tissue resource for the interpretation of genetic variation associated with complex disease. We provide openly available summary statistics of cis-eQTLs for all 44 tissues on the GTEx Portal (http://gtexportal.org) and all raw data in dbGaP (phs000424.v6.p1).

### Single-tissue cis-eQTL discovery

cis-eQTLs, or associations between local genetic variation and gene expression (≤ 1 Mb from the transcription start site, TSS), were identified using genotype and RNA-seq data generated from 44 tissues (N = 70–361 samples per tissue) using a linear model (FastQTL)^13^ (Fig. 1a,b). Within each tissue, we identified a median of 2,866 genes with cis-eQTLs at a 5% FDR (hereafter referred to as eGenes). In total, we found 159,760 cis-eQTLs for 20,175 genes, representing 82.6% of all genes tested in GTEx and 78.3% of all annotated autosomal lincRNA and protein coding genes^14^. For autosomal protein-coding genes alone, we identified 16,605 eGenes representing 90.2% of all expressed protein-coding genes in GTEx and 88% of all annotated protein-coding genes (Fig. 1c). For genes without an eQTL in any tissue, we observed less selective constraint as well as enrichment of functions related to transcriptional regulation, environmental response, and cellular differentiation, indicating that biological context influences the discovery of eQTLs for these genes (Extended Data Fig. 1). eGene discovery increased linearly with sample size with no evidence of saturation at the full sample size for each tissue, suggesting that all genes may ultimately be shown to be influenced by regulatory variation (Extended Data Fig. 2).

**Figure 1.**
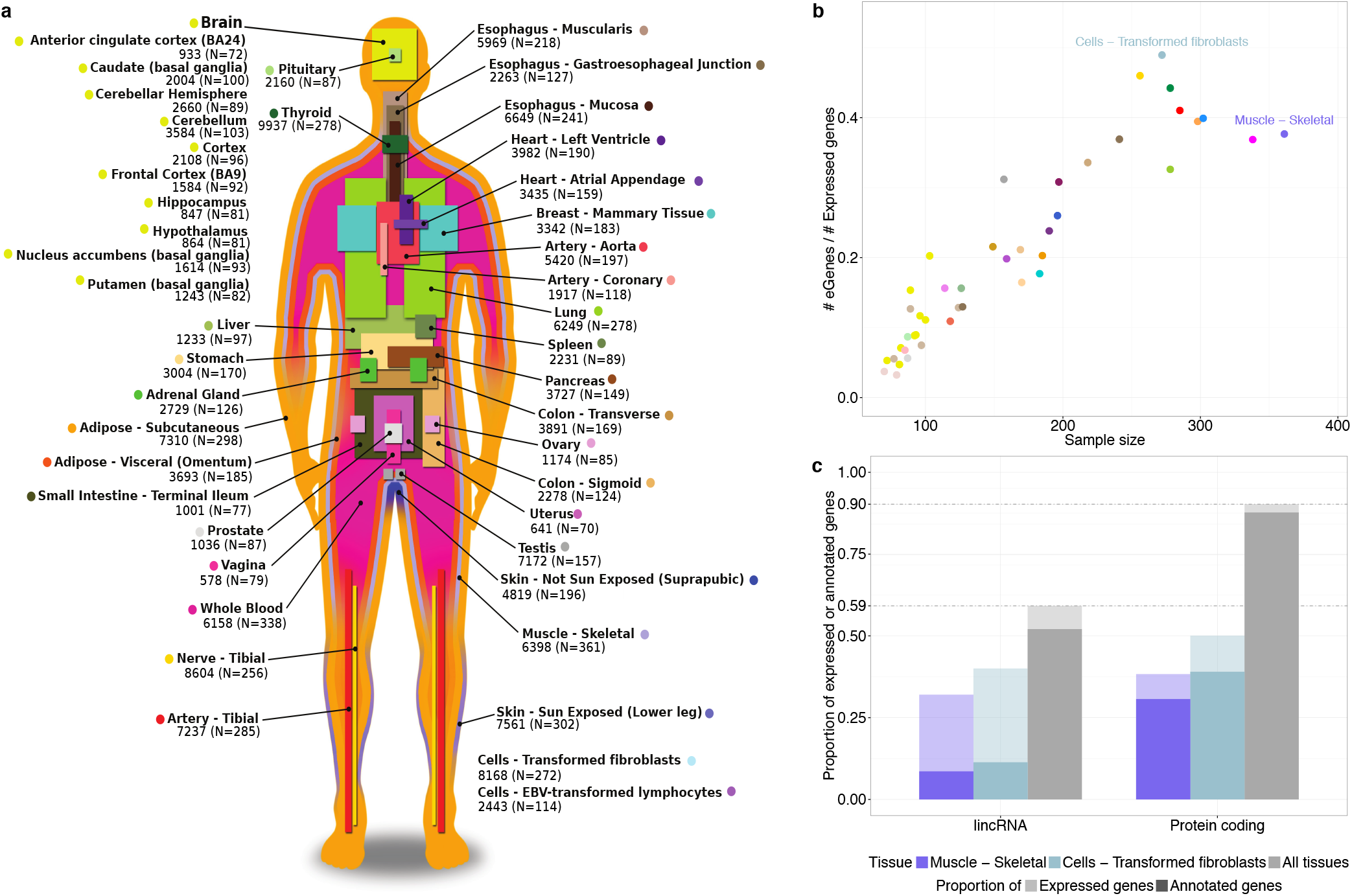
Sample size and eGene discovery in the GTEx V6p study. (a) Illustration of the 44 tissues and cell lines included in the GTEx V6p project with the associated number of eGenes and sample sizes. (b) The proportion of expressed genes discovered as eGenes versus sample size. Cells – Transformed fibroblasts are highlighted as the tissue with the highest proportion. Muscle – Skeletal has the largest sample size. (c) Fraction of genes that are eGenes across all tissues by transcript class. As in (b), Cells – Transformed fibroblasts and Muscle – Skeletal are shown as a reference. Annotated genes are all known human genes for each transcript class as curated in GENCODE v19.

We also identified conditionally independent regulatory variants for each eGene (secondary cis-eQTLs) using forward-backward stepwise regression separately in each tissue. This approach revealed an additional 22,099 cis-eQTLs across the 44 tissues, with 36.7% of protein-coding genes and 12.5% of lincRNAs having multiple, conditionally independent cis-eQTLs in at least one tissue (Extended Data Fig. 3).

The large sampling of tissues allowed us to develop a comprehensive view of the sharing of cis-eQTLs across tissues in the human body. We tested the replication of cis-eQTLs using the *π*_1_ statistic^15^ for all tissue pairs (Fig. 2a). We observed patterns of sharing that reflected previously identified relationships between tissues^1^. For example, we found a high degree of sharing between brain tissues (mean *π*_1_ of 0.864), arterial tissues (mean *π*_1_ of 0.854), and skeletal muscle and heart tissues (mean *π*_1_ of 0.819). The mean *π*_1_ sharing across all tissue pairs was 0.727 ranging from 0.354 to 0.981. Since individuals in the GTEx dataset contribute samples for multiple tissues, we investigated the effect of this grouping on sharing estimates by calculating *π*_1_ for tissues subsampled to have complete sharing among individuals (Extended Data Fig. 4). These sharing estimates correlated with estimates from variable levels of individual overlap between tissues (Spearman *ρ* = 0.53, *P <* 2.2 × 10^−16^). Furthermore, in the full dataset for each tissue, we observed that even for very strong shared associations (*P* < 10^−10^ in each tissue), roughly 10% exhibited different single top gene associations across tissues, indicating that the interpretation of the regulatory effect of these variants can still be tissue-dependent (Fig. 2b).

**Figure 2.**
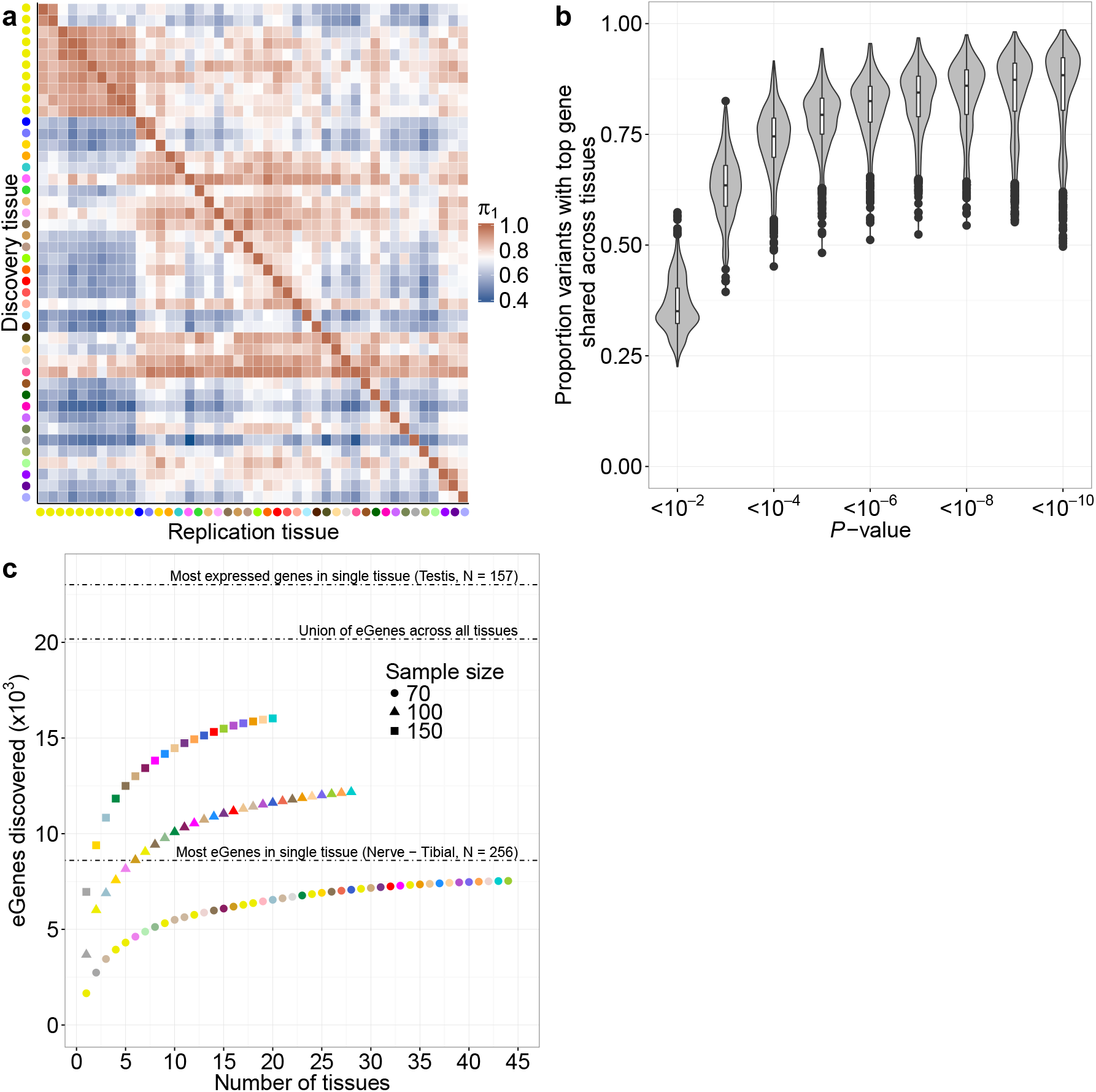
Single-tissue eQTL discovery across tissues. (a) Replication of eQTLs between tissues. Pairwise *π*_1_ statistics are reported for single-tissue eQTL discoveries in each tissue. Higher *π*_1_ values indicate an increased replication of eQTLs. Tissues are grouped using hierarchical clustering on rows and columns separately with a distance metric of 1 – *ρ*, where *ρ* is the Spearman correlation of *π*_1_ values. *π*_1_ is only calculated when the gene is expressed and testable in the replication tissue. (b) Proportion of variants with top associated protein-coding gene preserved between tissues shown for varying nominal association thresholds. (c) eGene discovery as a function of sample size and number of tissues assayed. Each tissue was subsampled to 70, 100, and 150 individuals and a greedy algorithm was used to assess sequential combinations of tissues that maximize the total number of unique eGenes discovered.

To quantify the impact of sample size and number of tissues studied on cis-eQTL discovery, we first compared eGene discovery across a range of sample sizes and tissues (Fig. 2c). The discovery of new eGenes was most influenced by sample size. However, a diverse sampling of tissues also improved eGene discovery. At its full sample size of 256 individuals, tibial nerve had the most eGenes of any tissue at 8,604, yet 9,394 unique eGenes were found for the top two tissues at a subsample size of 150 individuals. Cerebellum, Testis, Nerve – Tibial, and Thyroid were among the most effective tissues in increasing the total number of unique eGene discoveries. We next tested how sample size influenced patterns of cis-eQTL sharing across tissues. We observed that cis-eQTLs discovered in GTEx tissues with large sample sizes were less likely to be shared in other tissues, indicating that weaker associations identified in deeply sampled tissues remain difficult to replicate due to their smaller and possibly tissue-specific effects (Fig. 3; Extended Data Fig. 5).

**Figure 3.**
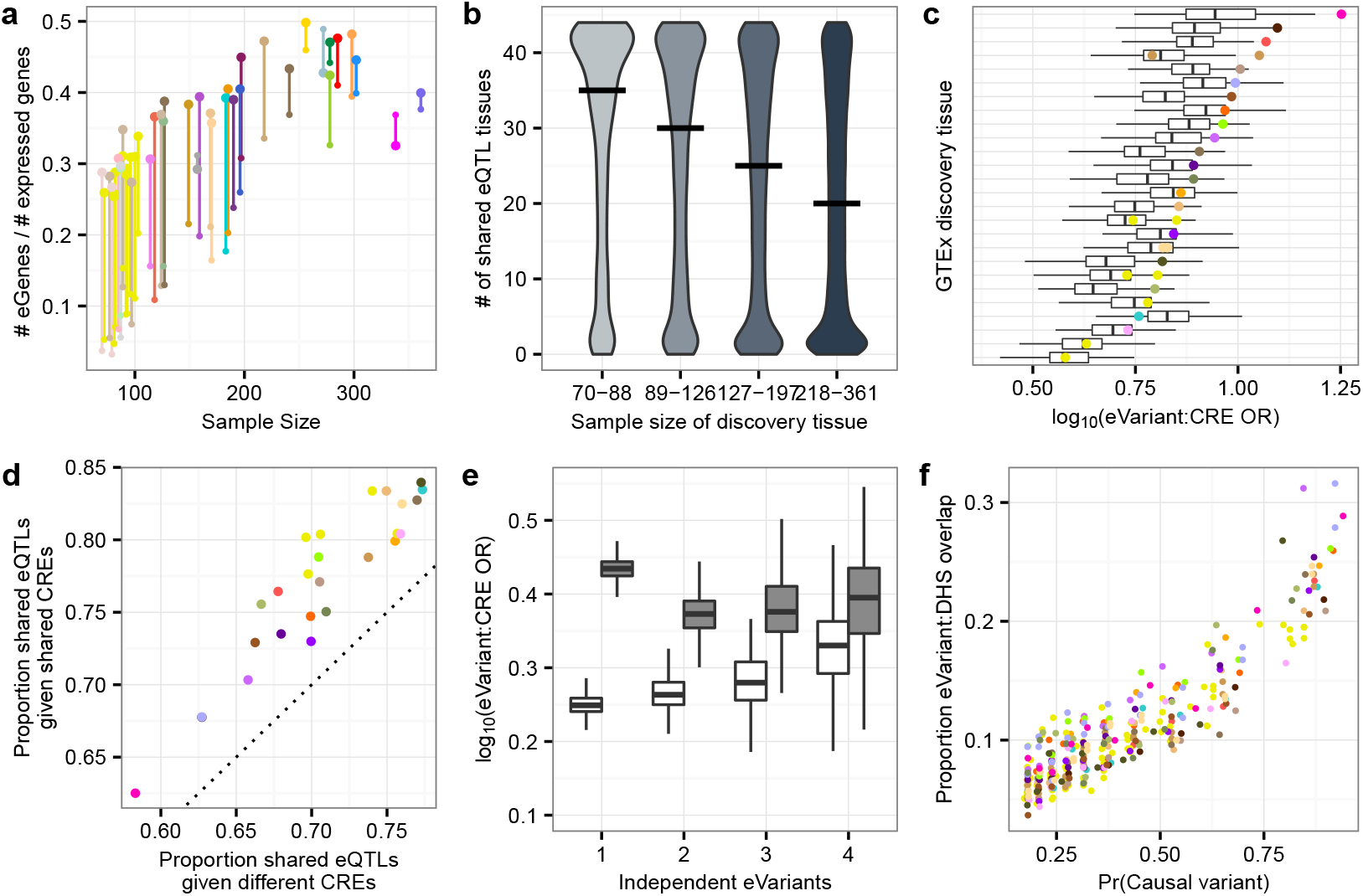
Multi-tissue eQTL discovery and genomic context. (a) The proportion of expressed genes for which eGenes are discovered in single tissues (5% FDR; small dots) and the multi-tissue meta-analysis (m-value ≥ 0.9; large dots), stratified by the sample size of individual tissues. In the meta-analysis, eQTL discoveries are made using METASOFT to identify tissues where the posterior probability a given eQTL effect exists (i.e. the tissue's m-value) is ≥ 0.9. (b) The number of tissues in which a given eQTL is shared as a function of tissue sample size. For each tissue, we calculated the degree of sharing (i.e. the number of tissues with m-value ≥ 0.9) for all eQTLs identified in that tissue at a 5% FDR. Tissues were then binned into quartiles based on sample size. The median number of shared tissues is plotted for each quartile as a horizontal black line. (c) Enrichment of eVariants in cis-regulatory elements (CREs) across 128 NIH Epigenomics Roadmaps cell types is depicted for each GTEx discovery tissue. Stronger enrichment was observed in matched tissues (colored dots) compared to unmatched tissues (boxplots). (d) Proportion of eQTLs that are shared between two tissues (m-value in both tissues ≥ 0.9) if the eVariant overlaps the same Roadmap annotation in both tissues (y-axis) or different annotations (x-axis). Points represent the mean of pairwise comparisons between all tissues, colored by the discovery tissue. (e) Enrichment of eVariants in tissue-matched enhancers (white) and promoters (grey) for the first four conditionally independent eQTLs discovered for each eGene (x-axis, sorted by discovery order). (f) Proportion of eVariants overlapping tissue-matched DNAse I hypersensitive sites as a function of the probability that a variant is causal. Points are colored by the eQTL discovery tissue.

### Multi-tissue cis-eQTL discovery

Multi-tissue cis-eQTL analyses have been shown to increase power while explicitly modeling sharing patterns across tissues^16–18^. We performed a meta-analysis across all 44 tissues using METASOFT^19^ and identified between 4,538 and 9,327 eGenes (m-value ≥ 0.9) per tissue. On average, each cis-eQTL effect was shared across 15 tissues. The advantage of meta-analysis was most apparent for individual tissues with smaller sample sizes (Fig. 3a), most notably for the 10 sampled brain regions. For example, in the hippocampus (N = 81), the number of single-tissue eGenes is 847 whereas the number of eGenes detected through meta-analysis is 4,636. eGenes identified by meta-analysis were more likely to be significant in single tissue analyses at larger sample sizes (Extended Data Fig. 6). Our meta-analysis approach demonstrates that sharing of cis-eQTL effects across multiple tissues can improve discovery in specialized or difficult-to-access tissues.

To ensure these findings did not depend on the modeling assumptions of METASOFT, we analyzed the FastQTL *P*-values for all genes and all tissues with TreeQTL, a hierarchical multiple comparison procedure, that controls the FDR of eGene discoveries across tissues^20^. This procedure identified 19,610 eGenes, 565 fewer eGenes than with the single-tissue analysis. While more conservative overall than the tissue-by-tissue analysis, we observed an increase in the number of eGenes detected in the tissues with the smallest sample sizes, as well as an increase in the average number of tissues in which an eGene is detected (from 7.9 for single-tissue analysis to 8.5; Extended Data Fig. 7).

Modeling of cis-eQTL sharing across tissues using METASOFT showed a bimodal pattern with increasing tissue-specificity for tissues with larger sample sizes (Fig. 3b). Increased tissue-specificity likely emerges from differences in discovery power and effect sizes across tissues. It also suggests that deep sampling diminishes the gains of meta-analysis, instead benefiting identification of more tissue-specific effects. The bimodal pattern of sharing was further supported by three different methods: simple overlap of the single-tissue results, the hierarchical procedure of TreeQTL, and an empirical Bayes model^18^ (Extended Data Fig. 8).

### Genomic features of cis-eQTLs

To characterize the genomic properties of cis-eQTLs, we annotated the associated variants (hereafter referred to as eVariants) with chromatin state predictions from 128 cell types sampled by the Roadmap Epigenomics Consortium, including 26 tissues that match GTEx tissues^21^. eVariants were enriched in predicted promoter and enhancer states across a broad range of tissues and exhibited significantly greater enrichment in promoters and enhancers from their matched tissues (linear model controlling for discovery cell type, *P* < 5.7 × 10^−10^), illustrating consistent patterns of cell type specificity for both cis-regulatory elements (CREs) and cis-eQTLs (Fig. 3c,e). Furthermore, cis-eQTLs were more likely to be active across pairs of tissues if the eVariant overlapped the same chromatin state in both tissues (paired Wilcoxon signed rank test, *P* < 2.2 × 10^−16^, Fig. 3d).

Compared to primary eVariants, secondary eVariants were located on average further away from the TSS (median distance 50.1 kb from the TSS versus 28.9 kb, Wilcoxon rank sum test, P < 2.2 × 10^−16^; Extended Data Fig. 9a) and exhibited less tissue sharing than primary eQTLs (Wilcoxon rank sum test, *P* < 2.2 × 10^−16^; Extended Data Fig. 9b). Both primary and secondary eVariants were enriched for promoter Hi-C contacts compared to background variant-TSS pairs (Wilcoxon rank sum test, *P* < 2.2 × 10^−16^; Extended Data Fig. 9c). This observation suggests that, despite their genomic distance from the TSS, many primary and secondary eVariants remain in close physical contact with their target gene promoters via chromatin looping interactions. Although primary eVariants are significantly more enriched in promoters than enhancers (Wilcoxon rank sum test, *P <* 2.2 × 10^−16^), secondary eVariants show greater enrichment in enhancers, consistent with their increasing distance from the TSS and tissue-specific activity (Wilcoxon rank sum test, *P* < 2.2 × 10^−16^; Fig. 3e; Extended Data Fig. 9c). This result underscores the importance of analyzing eQTLs beyond the primary association to discover regulatory variants in enhancers, which are known to be particularly relevant for disease associations^22–24^.

Integration of genomic annotations in eQTL testing has been demonstrated to improve power^6^, ^25–27^. We applied a Bayesian hierarchical model incorporating variant-level genomic annotations for eQTL discovery in 26 tissues with cell-type matched annotations from the Epigenomics Roadmap^28^ (Wen, X. submitted). Distance to the TSS and promoter and enhancer annotations improved our ability to discover eQTLs (Extended Data Fig. 10a). Using these annotations increased the total number of eGene discoveries by an average of 43% (1,200 genes) across tissues (Extended Data Fig. 10b).

### Fine-mapping eQTL variants

To identify likely causal variants underlying eQTLs, we applied two computational fine-mapping strategies. First, we identified 90% credible sets for each eGene in each tissue using CAVIAR^29^, a probabilistic method that utilizes the observed marginal test statistics and LD structure to detect variant sets that may harbor more than one causal variant^29^. Across all tissues, the mean credible set size was 29 variants (per tissue means ranged from 25 to 31). Credible set size decreased with increasing discovery tissue sample size. The addition of 100 samples reduced credible set size by an average of one variant indicating that large sample sizes are required to identify causal variants using association strength alone (Extended Data Fig. 11a). As expected, credible sets overlapped across tissues more extensively for tissue-shared eQTLs compared to tissue-specific eQTLs (Extended Data Fig. 11b).

We estimated the probability that each eVariant is a causal variant using CaVEMaN, a non-parametric sampling-based approach that accounts for noise in expression measurements and linkage structure (Brown et al. in preparation). Across tissues, we estimated that between 3.5%-11.7% of primary eVariants are causal (probability ≥ 0.8; Extended Data Fig. 12). For predicted causal variants, the same variant is predicted as causal for 13.3% to 32.6% of variants at the same probability threshold in separate tissues where an eGene is also identified. However, the replication rate *π*_1_ was considerably higher (59.6%-93.5%), demonstrating the difficulties in fine mapping variants even when the LD structure is expected to be preserved across tissues. Consistent with predicted causal variants being functional regulatory variants (as opposed to LD proxies), 24.3% of eVariants with causal probabilities in the top 10th percentile (P > 0.77) overlapped open chromatin regions compared to 11.2% of all eVariants and 6.6% of eVariants in the lowest 10th percentile (0.027 < P < 0.19; Fig. 3f).

### cis-eQTL effect sizes

To determine the effect sizes of eQTLs discovered in GTEx, we used an additive model of eQTL alleles on total gene expression, allowing for biologically meaningful interpretation of effect sizes as an allelic fold change between the two eQTL alleles (see Methods; Mohammadi et al. in preparation). 17.4% of eGenes had eQTLs with median effect sizes of ≥ 2-fold across tissues (Fig. 4a). As expected, mean effect sizes per tissue were influenced by sample size (Extended Data Fig. 13). When stratifying each gene by the number of tissues that it is expressed in, we observed a decrease in the average effect size per gene indicating that genes expressed in multiple tissues are less likely to have eQTLs with large regulatory effects (Spearman **ρ** = -0.29, *P <* 2.2 × 10^−16^, Fig. 4b). Supporting this observation, tissue-shared eQTLs had significantly smaller effect sizes than tissue-shared eQTLs matched for significance level (Wilcoxon rank sum test, P < 2.2 × 10^−16^, Fig. 4c; Extended Data Fig. 13).

**Figure 4.**
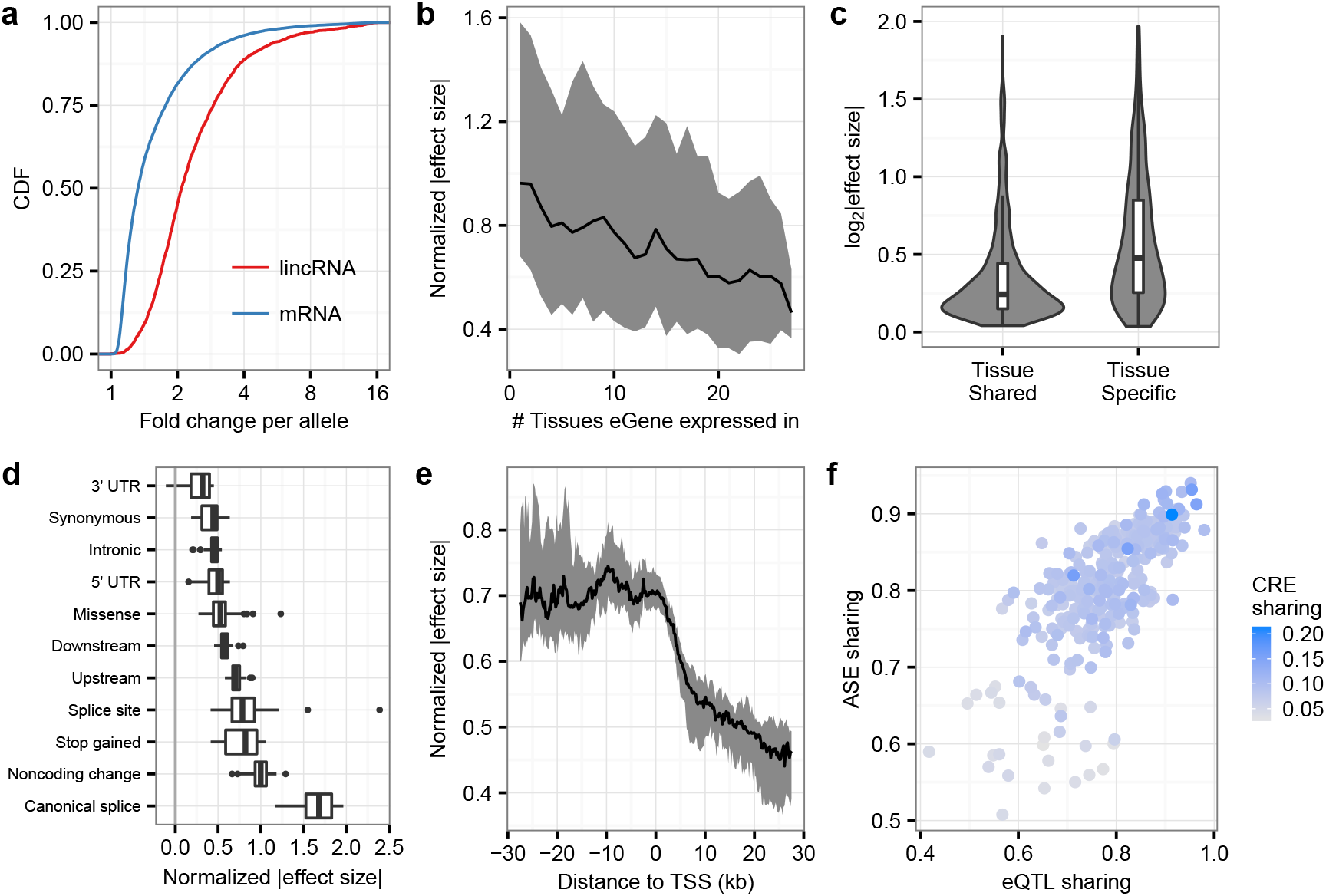
ASE and the epigenomic context of cis-eQTLs across tissues. (a) For each autosomal protein-coding and lincRNA eGene, the median effect size was computed across all tissues with eVariants for that eGene. The empirical CDF of these median effect sizes is depicted. (b) Median (line) and interquartile range (ribbon) of absolute eQTL effect size, corrected for median expression level across tissues and the minor allele frequency of the eVariant, as a function of the number of tissues the eGene is expressed in. (c) Comparison of effect sizes between q-value-matched tissue-shared eQTLs (m-value > 0.9 in at least 35 tissues) and tissue-specific eQTLs (m-value ≥ 0.9 in only the discovery tissue). (d) Normalized absolute eQTL effect size for each top eVariant, for each eVariant annotation. Normalized effect sizes were estimated by correcting for eVariant minor allele frequency and cross tissue effect size differences. (e) Normalized (as in d) eQTL effect size depicted in 200bp bins, relative to the eGene TSS. Bin medians and interquartile ranges plotted as lines and ribbons, respectively. (f) Pairwise tissue sharing of ASE effects for genes with bimodal ASE effects (proportion with same ASE mode; y-axis) is correlated with pairwise eQTL sharing *(π*_1_, x-axis), and the fraction of eVariants overlapping the same Roadmap annotation in both tissues.

We assessed whether variants with distinct functional annotations had different average effects on gene expression using the large number of eQTLs we discovered. eVariants at canonical splice sites exhibited the strongest effects, followed by variants in noncoding transcripts (Fig. 4d). Variants in the 3' UTR had the weakest effect, significantly weaker than those in 5' UTRs (Wilcoxon rank sum test, *P <* 4.81 × 10^−10^). Missense variants had a significantly stronger effect on gene expression than synonymous variants (Wilcoxon rank sum test, *P <* 8.65 × 10^−5^). Analysis of eQTL effect sizes around the TSS demonstrated that upstream variants had a stronger effect on gene expression than downstream variants (Wilcoxon rank sum test, *P <* 1.94 × 10^−15^; Fig. 4e), an effect that seems to persist through the gene body and beyond. These results suggest that eVariants likely to affect transcription have stronger effects on gene expression levels than variants likely to impact post-transcriptional regulation of mRNA levels.

### Allele-specific expression (ASE)

The impact of a regulatory variant on expression may be estimated from either total expression or allele specific expression (ASE) estimates. We measured ASE^30^ at over 135 million sites across tissues and individuals, with a median of over 10,000 genes quantified per donor (Extended Data Fig. 14a-f). In total, 63.5% of all protein-coding genes could be tested for ASE in at least one individual and tissue with 62.6% having ASE data from multiple individuals in at least one tissue. 87.9% of testable genes had significant allelic imbalance in at least one individual (binomial test, FDR < 0.05), demonstrating an abundance of cis-linked regulatory effects. Across individuals, a median of 1,963 genes had significant allelic imbalance in at least one tissue, with a median of 570 genes where the individual was not heterozygous for a top eQTL. We independently estimated the effects of the primary eVariant for each eGene in each tissue using both allele-specific and total gene expression measurements (see Methods). Effect size estimates from both approaches are highly consistent with an average Spearman correlation of 0.84 (std. dev. = 2%; Extended Data Fig. 15) and an average ratio of ASE effect size to eQTL effect size of 98.5% (std. dev. = 1%). This observation confirms that cis-eQTLs and ASE capture the same biological phenomenon.

We modeled allelic expression in genes across different tissues of each individual in order to capture tissue-specificity of regulatory variant function. Over 17% of genes exhibit allelic expression patterns that differed across tissues in at least one individual. Patterns of ASE sharing in these genes were used to cluster tissues independently of total gene expression levels, which may be more susceptible to shared environmental influences, and without the strong dependency with sample size that complicates analyses of eQTL sharing (Extended Data Fig. 14g; Fig. 2a). Indeed, pairwise ASE sharing was highly correlated with pairwise eQTL sharing (Spearman *ρ* = 0.70, *P* < 2.2 × 10^−16^). Moreover, both pairwise ASE and eQTL sharing are correlated with pairwise tissue sharing of eVariant CRE annotation (Spearman *ρ* > 0.29, *P* < 2.6 × 10^−7^; Fig. 4f).

### eQTLs and GWAS

The expanded GTEx resource provides a unique opportunity to interpret GWAS associations for a wide range of complex traits and diseases. The increased diversity of tissue sampling has resulted in more identified tissue-specific eQTLs. Indeed, the degree of tissue sharing of an eQTL is associated with several indicators of phenotypic impact. eGenes shared across many tissues harbor fewer protein-coding loss-of-function (LoF) variants curated in the ExAC database^31^ (Fig. 5a), consistent with purifying selection removing large effect regulatory variants that involve many tissues. Tissue-shared eGenes were also less likely to be annotated disease genes compared to tissue-specific eGenes (Fisher’s exact test, nominal *P <* 10^−6^ for GWAS, OMIM, and LoF intolerant gene sets; Fig. 5a, Extended Data Fig. 16), highlighting that the cell-type specific mechanisms underlying complex genetic diseases may be elucidated only through broad tissue sampling.

**Figure 5.**
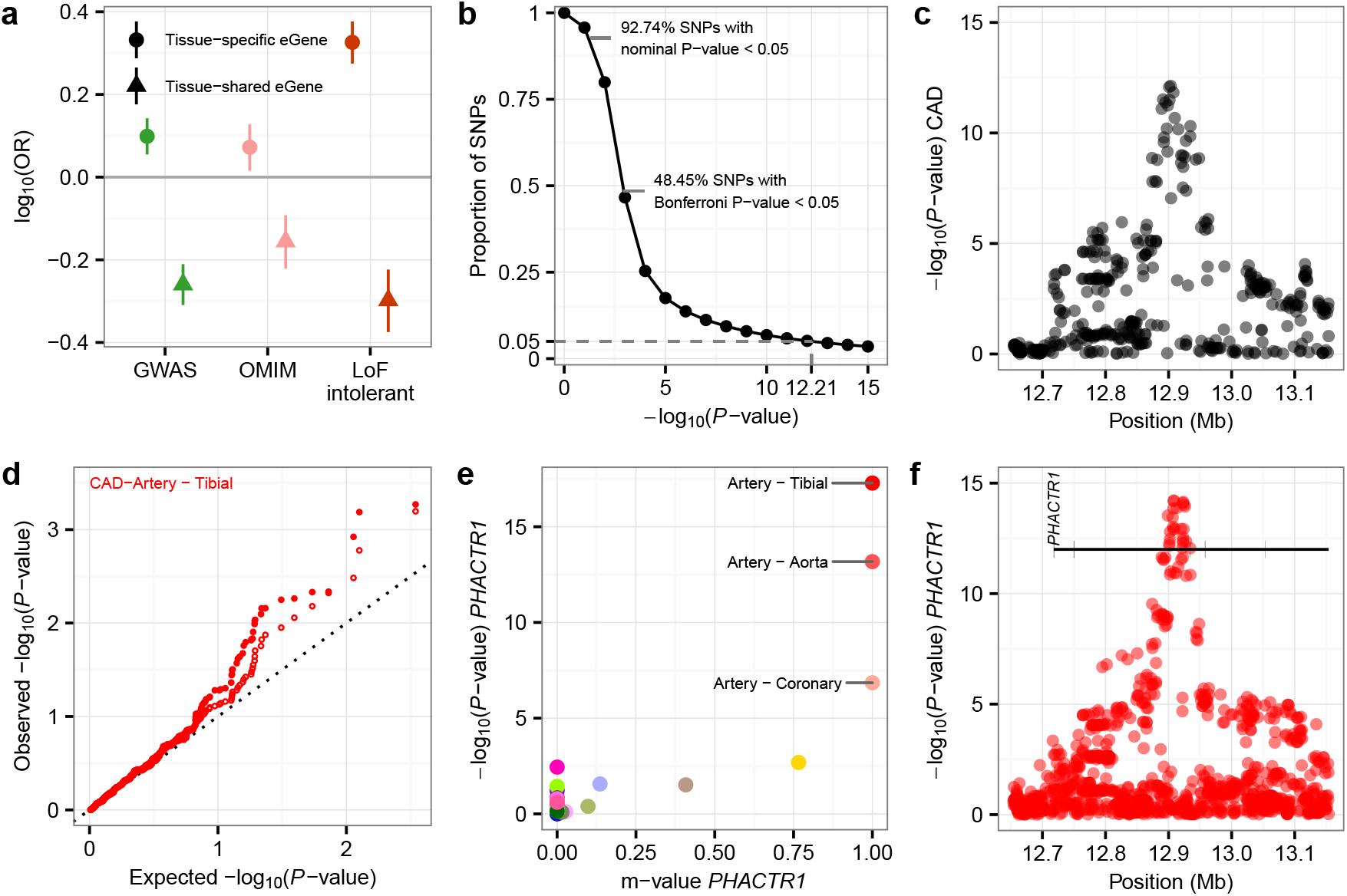
Intersection of cis-eQTLs with GWAS. (a) Enrichment of tissue-specific and tissue-shared eGenes in disease and loss of function mutation intolerant genes. eGenes were defined in each tissue using METASOFT (m-value ≥ 0.9). Tissue-specific and shared eGenes were defined as eGenes in the bottom and top 10% of the distribution of proportion of tissues with an eQTL effect, respectively. Points represent the log odds ratio for enrichment of the eGene category in each gene list. Bars represent 95% confidence intervals (Fisher's exact test). (b) Proportion of eQTLs discovered as a function of *P*-value cutoffs. Nearly 93% of all SNPs passed a nominal significance threshold of 0.05. More than 48% of all SNPs passed a Bonferroni threshold defined as the nominal threshold divided by the number of tissues (44). To control the type I error rate at 5%, a stringent cutoff of 10^−12^ is needed. (c) CAD association significance (y-axis) for all SNPs within 250 kb of the sentinel SNP (x-axis). (d) Quantile-quantile plot for CAD GWAS associations. Observed GWAS *P*-values (y-axis) plotted as a function of expected *P*-values (x-axis), for the top 1,000 eQTLs (closed circles) and MAF and distance matched SNPs (open circles). (e) For each tissue, the METASOFT m-value (x-axis) of the lead CAD SNP is plotted against the single-tissue eQTL association significance (y-axis). Points are colored by tissue. (f) eQTL association significance (y-axis) for *PHACTR1* for all SNPs within 250 kb of the sentinel CAD SNP (x-axis).

This broad sampling affects the interpretation of eQTL data in the context of GWAS variants. We observed that 92.7% of all common variants assayed by GTEx are nominally associated with the expression of one or more genes in one or more tissues (P < 0.05) and nearly 50% are significant when performing a Bonferroni correction based on the number of tissues tested (Fig. 5b). Given the ubiquity of eQTL associations, caution is warranted when using eQTL data to interpret the function of candidate variants without assessing whether GWAS and eQTL association signals are likely driven by the same causal variant by colocalization approaches that examine local LD and trait summary statistics^22,32–34^.

To illustrate the utility of GTEx for the interpretation of disease-associated variation, we applied GTEx to the *PHACTR1* locus which is associated with a range of complex traits, including myocardial infarction (MI)^35^, coronary artery disease (CAD)^36–38^, cervical artery dissection^39^, and migraines^40^ (Fig. 5c). Notably, the CAD and MI risk allele (G) at rs9349379 is protective for cervical artery dissection and migraines. Initial targeted analyses^41^ demonstrated that the CAD risk allele (G) at rs9349379 is associated with decreased expression of *PHACTR1* in coronary arteries.

To investigate the mechanism and tissue of action of this pleiotropic SNP, we characterized the effect of rs9349379 across the 44 GTEx tissues. rs9349379G was strongly associated with decreased *PHACTR1* expression (Fig. 5e; meta-analysis P < 2.2 × 10^−16^), with a tissue-specific eQTL effect observed only in aorta, coronary, and tibial arteries (Fig. 5e; m-value ≥ 0.9), where the risk allele expression is 72%, 57% and 65% of the protective allele expression, respectively. *PHACTR1*, *TBC1D7*, and the nearby noncoding RNA, *RP1-257A7.5*, were the only genes within 1 Mb associated with genotype at rs9349379 in any tissue. Notably, the tissue specificity of the eQTL effect was not mirrored in the tissue specificity of *PHACTR1* gene expression (Extended Data Fig. 17). Colocalization analysis in arterial tissues indicated that rs9349379 is likely the variant responsible for both the GWAS and the eQTL signal in the locus (Fig. 5f; RTC = 1, eCAVIAR = 0.95)^34,42^. Applying the PrediXcan method^12^ to the BioVU repository^43^, we found that genetically predicted decreased *PHACTR1* expression in coronary and aorta arteries was associated with tachycardia (meta-analysis P < 10^−6^), whereas genetically predicted increased *PHACTR1* expression was associated with migraines (*P* = 1.2 × 10^−7^). *PHACTR1* is the sole gene in the locus that was implicated by PrediXcan in BioVU, using arterial tissues, for either trait. These results suggest that the pleiotropic effects of rs9349379 are driven by a consistent, tissue-specific molecular phenotype that causes diverse downstream consequences.

## Discussion

The most immediate effects of functional genetic variation are on molecular phenotypes. Combining trait and disease associated variants with molecular QTL data has been a successful strategy for resolving causal genes and tissues^44^. In particular, these approaches have provided key information on human-specific traits and therapeutic interventions^11,45,46^. While the pilot phase of the GTEx project identified cis-eQTLs in nine tissues, the GTEx V6p collection has been expanded to 44 tissues providing a wealth of additional cis-eQTL discoveries. These data facilitate both systematic and targeted interpretation of the functional consequences of genetic variants across a range of biological contexts.

We found a pervasive effect of common regulatory variation on the vast majority of human genes with a sizable proportion of genes having multiple independent loci associated with their expression levels. By combining cis-eQTL data across tissues, we demonstrated that GTEx V6p data may be used to enable cis-eQTL discovery in tissues with limited sample sizes. Many of the largest, primary effects are shared across tissues. Additionally, we observed that both secondary cis-eQTLs and cis-eQTLs from deeply sampled tissues exhibit more tissue-specificity. cis-eQTLs are enriched in both tissue-specific enhancers and promoters and patterns of regulatory element overlap are predictive of tissue sharing for cis-eQTLs. Secondary cis-eQTLs were as enriched as primary cis-eQTLs for Hi-C contacts suggesting a direct effect on gene expression facilitated by chromosome looping and local nuclear organization. Furthermore, we demonstrated that tissue-specific genes and eQTLs have larger effect sizes, and we have presented a large resource of allelic expression data that demonstrates correlated estimates of tissue-sharing and effect size estimates with eQTLs. Overall, these observations illustrate the systemic effects of regulatory variants and inform eQTL study design by highlighting the unique contributions of tissue-specific eQTLs that can only be identified through broad tissue sampling.

The wealth of cis-eQTLs identified in this study bears important implications for GWAS interpretation. We demonstrated that 92.7% of variants tested in our study have a nominally significant association with expression (*P*-value < 0.05) and that approximately 10% of eVariants may change their top associated gene when tested in another tissue. Given the abundance of associations for any variant, care must be taken in using cis-eQTL data to propose novel biological mechanisms for disease-associated variants. GWAS variants are enriched among tissue-specific cis-eQTLs, highlighting the necessity for sampling of diverse tissues. The wealth of associations in GTEx may further aid in selecting candidate causal tissues where multiple GWAS signals for specific traits are enriched. Within these targeted tissues, colocalization strategies that combine locus-specific trait and expression association information are required to understand the underlying biological mechanism. We showed that a combination of these approaches may be used to interpret a GWAS signal of relevance to coronary artery disease, and we revealed novel tissue-specific biology identified through the analysis of GTEx V6p data.

Together, cis-eQTL data in GTEx V6p provide the most comprehensive characterization of the local effects of regulatory variation to date. We expect that these data will be of considerable utility for the interpretation of gene regulatory mechanisms, human evolution, and complex trait and disease biology.

## Online Methods

### Sample procurement

The GTEx V6p eQTL analysis freeze represents 44 distinct tissue sites collected from 449 postmortem donors representing a total of 7,051 tissues. All human subjects were deceased donors. Informed consent was obtained for all donors via next-of-kin consent to permit the collection and banking of de-identified tissue samples for scientific research. Complete descriptions of the donor enrollment and consent process, as well as biospecimen procurement, methods, sample fixation and histopathological review procedures were previously described^1,47^. Briefly, whole blood was collected from each donor, along with fresh skin samples, for DNA genotyping, RNA expression and culturing of lymphoblastoid and fibroblast cells, and shipped overnight to the GTEx Laboratory Data Analysis and Coordination Center (LDACC) at the Broad Institute. Two adjacent aliquots were then prepared from each sampled tissue and preserved in PAXgene tissue kits. One of each paired sample was embedded in paraffin (PFPE) for histopathological review, the second was shipped to the LDACC for processing and molecular analysis. Brains were collected from approximately 1/3rd of the donors, and were shipped on ice to the brain bank at the University of Miami, where 11 brain sub-regions were sampled and flash frozen. These samples were also shipped to the LDACC at the Broad Institute for processing and analysis.

All DNA genotyping was performed on blood-derived DNA samples, unless unavailable, in which case a tissue-derived DNA sample was substituted. RNA was extracted from all tissues, but quality varied^1^. RNA sequencing was performed on all samples with a RIN score of 5.7 or higher and with at least 500ng of total RNA. Nucleic acid isolation protocols, and sample QC metrics applied, are as described in^1^.

### Data production

RNA was isolated from a total of 9,547 postmortem samples from 54 tissue types from up to 550 individuals. 44 tissues were sampled from at least 70 individuals: 31 solid-organ tissues, 10 brain subregions with two duplicate regions (cortex and cerebellum), whole blood, and two cell lines derived from donor blood and skin samples. Each tissue had a different number of unique samples. Non-strand specific, polyA+ selected RNA-seq libraries were generated using the Illumina TruSeq protocol. Libraries were sequenced to a median depth of 78 million 76-bp paired end reads. RNA-seq reads were aligned to the human genome (hg19/GRCh37) using TopHat^48^ (v1.4) based on GENCODE v19 annotations^14^. This annotation is available on the GTEx Portal (gencode.v19.genes.v6p model.patched contigs.gtf.gz). Gene-level expression was estimated as reads per kilobase of transcript per million mapped reads (RPKM) using RNA-SeQC on uniquely mapped, properly paired reads fully contained with exon boundaries and with alignment distances ≤ 6. Samples with less than 10 million mapped reads or with outlier expression measurements based on the D-statistic were removed^49^.

DNA isolated from blood was used for genotyping. 450 individuals were genotyped using Illumina Human Omni 2.5M and 5M Beadchips. Genotypes were phased and imputed with SHAPEIT2^50^ and IMPUTE2^51^, respectively, using multi-ethnic panel reference from 1000 Genomes Project Phase 1 v3^52^. Variants were excluded from analysis if they: (1) had a call rate < 95%; (2) had minor allele frequencies < 1%; (3) deviated from Hardy-Weinberg Equilibrium (*P* < 10^−6^); or (4) had an imputation info score less than 0.4.

### cis-eQTL mapping

We conducted cis-eQTL mapping within the 44 tissues with at least 70 samples each. Only genes with ≥ 10 individuals with expression estimates > 0.1 RPKM and an aligned read count ≥ 6 within each tissue were considered significantly expressed and used for cis-eQTL mapping. Within each tissue, the distribution of RPKMs in each sample was quantile-transformed using the average empirical distribution observed across all samples. Expression measurements for each gene in each tissue were subsequently transformed to the quantiles of the standard normal distribution. The effects of unobserved confounding variables on gene expression were quantified with PEER^53^, run independently for each tissue. 15 PEER factors were identified for tissues with less than 150 samples; 30 for tissues with sample sizes between 150 and 250; and 35 for tissues with more than 250 tissues.

Within each tissue, cis-eQTLs were identified by linear regression, as implemented in FastQTL^13^, adjusting for PEER factors, gender, genotyping platform, and three genotype-based PCs. We restricted our search to variants within 1 Mb of the transcription start site of each gene and, in the tissue of analysis, minor allele frequencies ≥ 0.01 with the minor allele observed in at least 10 samples. Nominal *P*-values for each variant-gene pair were estimated using a two-tailed t-test. Significance of the most highly associated variant per gene was estimated by adaptive permutation with the setting `"--permute 1000 10000"`. These empirical *P*-values were subsequently corrected for multiple testing across genes using Storey’s q-value method^15^.

To identify the list of all significant variant-gene pairs associated with eGenes, a genome-wide empirical *P*-value threshold, *p*_*t*_, was defined as the empirical *P*-value of the gene closest to the 0.05 FDR threshold. *p*_*t*_ was then used to calculate a nominal *P*-value threshold for each gene based on the beta distribution model (from FastQTL) of the minimum *P*-value distribution *f*(*p*_min_) obtained from the permutations for the gene. Specifically, the nominal threshold was calculated as *F*^−1^(*p*_*t*_), where *F*^−1^ is the inverse cumulative distribution. For each gene, variants with a nominal *P*-value below the gene-level threshold were considered significant and included in the final list of variant-gene pairs.

### Multi-tissue cis-eQTL mapping

To increase sensitivity of cis-eQTL detection, in particular of cis-eQTLs with smaller effect sizes, we ran METASOFT^54^, a meta-analysis method, on all variant-gene pairs that were significant (FDR < 5%) in at least one of the 44 tissues based on the single-tissue results from FastQTL. The goal of this analysis was to gain power to discover additional tissues for a cis-eQTL. A random effects model in METASOFT (called RE2), designed to find loci with effects that may have heterogeneity between datasets/tissues (and assumes estimates are independent and consistent in effect direction) was used^19^. The posterior probability that an eQTL effect exists in a given tissue, or m-value^54^, was calculated for each variant-gene pair and tissue tested. A significance cutoff of m-value ≥ 0.9 was used to discover high-confidence cis-eQTLs.

We applied a separate hierarchical multiple testing correction method to identify multi-tissue eGenes. First, we constructed a P-value for each eGene across tissues using the Simes combination rule^55^ on the tissue-specific beta-approximation P-values provided by FastQTL. Storey’s q-value method^15^ was then used to identify eGenes that are active in any tissue. To identify the specific tissues in which these eGenes are regulated, we applied the Benjamini and Bogomolov procedure^56^ at the 0.05 level. This approach not only allowed us to control the FDR for the discovery of eGenes across tissues and the expected average proportion of false tissue discoveries across these eGenes, but also to gain power to detect eGenes in tissues with smaller sample sizes when there is evidence from other tissues supporting their regulation.

### Independent cis-eQTL mapping

#### Single-tissue analysis

Multiple independent signals for a given expression phenotype were identified by forward stepwise regression followed by a backwards selection step. The gene-level significance threshold was set to be the maximum beta-adjusted P-value (correcting for multiple-testing across the variants) over all eGenes in a given tissue. At each iteration, we performed a scan for cis-eQTLs using FastQTL, correcting for all previously discovered variants and all standard GTEx covariates. If the beta adjusted *P*-value for the lead variant was not significant at the gene-level threshold, the forward stage was complete and the procedure moved on to the backward stage. If this *P*-value was significant, the lead variant was added to the list of discovered cis-eQTLs as an independent signal and the forward step moves on to the next iteration. The backwards stage consisted of testing each variant separately, controlling for all other discovered variants. To do this, for each eVariant, we scanned for cis-eQTLs controlling for standard covariates and all other eVariants. If no variant was significant at the gene-level threshold the variant in question was dropped, otherwise the lead variant from this scan, which controls for all other signals found in the forward stage, was chosen as the variant that represents the signal best in the full model.

#### Multi-tissue analysis

We ran a modified version of forward stepwise regression to select an ordered list of independent variants associated with a given gene across all tissues types. In each step k, we identify variants associated with expression of each gene across tissues, and refer to these as the tier k variants. In each tier k, for each tissue, Matrix-eQTL was run independently for each gene that had a variant added to the model at every previous step 1..*k*–1 (all genes are assessed in tier 1). In each tier, any significant variants identified in tiers 1..*k*–1 are included as covariates. Significant tier k variants were assessed as follows. For each tissue, we obtained gene-level *P*-values for tier *k* via eigenMT^57^. Genome-wide significance of multiple independent variants per gene (in each tissue independently) was assessed via Benjamini-Hochberg (FDR < 0.05) for all gene-level *P*-values tested in tier k combined with all those tested in previous tiers^58^. To identify the cross-tissue tier k variant for a given gene, we selected the variant (out of all variants genome-wide significant for the gene in at least one tissue) with the smallest geometric mean *P*-value (across tissues). If no variant was genome-wide significant, no cross-tissue tier *k* variant was selected for that gene, and that gene will be estimated to have *k* – 1 total independent cross-tissue variants. If a particular tissue’s tier *j* genome-wide significant variant for a particular gene differed from the cross-tissue tier *j* variant for the same gene, the *P*-value of that tissue‘s tier *j* genome-wide significant variant was used in the Benjamini-Hochberg procedure. If a particular gene‘s cross-tissue variant for tier *k* does not meet genome-wide significance in all tissues in the tier (*k* + 1) step due to increased multiple testing, that gene will be conservatively considered to have (*k −* 1) independent cross-tissue variants.

### Allele-specific expression

#### Data generation

For each sample, allele-specific RNA-seq read counts were generated at all heterozygous SNPs with the GATK ASEReadCounter tool using default settings^30^. Only uniquely mapping reads with a base quality ≥ 10 at the SNP were counted, and only those SNPs with coverage of at least 8 reads were reported. Unless otherwise mentioned, SNPs that met any of the following criteria were flagged and removed from downstream analyses: (1) UCSC 50mer mappability of < 1, (2) simulation-based evidence of mapping bias^59^, (3) heterozygous genotype not supported by RNA-seq data across all samples for that subject (test adapted from Castel et al.^30^). Phasing between variants was determined using population phasing, and for some analyses was used to aggregate allelic counts across variants. Full ASE data is available through dbGAP.

#### Modeling patterns of ASE sharing across tissues

We used a beta-binomial mixture to model ASE across tissues, with each component corresponding to a distinct mode of allelic imbalance. The model was learned independently for each heterozygous coding SNP in each individual. Optimization was performed using five independent initial parameters values. The number of components in the mixture model, *K,* was selected using Bayesian Information Criterion (BIC). Variance of the BIC was estimated by bootstrapping and the most parsimonious model within one standard deviation of the global minimum BIC model was chosen as the optimal model.

Individuals with RNA-seq data from at least 20 tissues were included in the analysis *(N =* 131). The most highly expressed, coding, heterozygous SNP in each gene was selected. Genes with at least 30 reads in at least two tissues and at least one tissue with allelic imbalance (defined as P < 10^−3^ under a binomial null model) were included in the analysis. In total 207,943 SNPs spanning 13,030 genes were modeled using 1, 2, 3, and 4 modes of allelic imbalance. 2% of SNPs exhibited more than one pattern of allelic imbalance across tissues (*K*=2: 4219 cases, *K=*3: 64 cases, and *K*=4: 4 cases). These multimodal cases involved 2,226 genes across individuals. SNPs with bimodal (*K*=2) pattern of allelic expression were used to derive estimates of ASE tissue sharing. Tissues with less than 100 cases were excluded from analysis. Tissue similarity was measured as the proportion of times two tissues exhibit the same mode of allelic imbalance.

### Effect size estimation

#### cis-eQTL effect size

cis-eQTL effect size was defined as the ratio between the expression of the haplotype carrying the alternative eVariant allele to the one carrying the reference allele in log2 scale and was calculated using the method presented in (Mohammadi et al. in preparation). In short, the model assumes an additive model of expression in which the total expression of a gene in a given genotype group is the sum of the expression of the two haplotypes: *e*(genotype) = 2*e*_*r*_, *e*_*r*_ + *e*_*a*_, 2*e*_*a*_, for reference homozygotes, heterozygotes, and alternate homozygotes, respectively, where *e*_*r*_ is expression of the haplotype carrying the reference allele and *e*_*a*_, expression of the haplotype carrying the alternative allele is: *e*_*a*_ = *ke*_*r*_ where 0 < *k <* ∞.

cis-eQTL effect size is represented in log2 scale as *s* = log_2_ *k*, and is capped at 100-fold to avoid outliers *(|s| <* log_2_ 100). Expression counts were retrieved for all top eGenes in all tissues and PEER corrected. Data was log-transformed with one pseudo-count to stabilize the variance. The model was fit using non-linear least squares to derive maximum likelihood estimates of the model parameters *k* and *e*_*r*_. A similar maximum likelihood approach with additive effects and multiplicative errors (prior to log transformation)^60^ was compared in several tissues to the effect size estimates reported here, exhibiting rank correlation 0.98. Confidence intervals for the effect sizes were derived using bias corrected and accelerated (BCa) bootstrap with 100 samples.

For all analyses in a given tissue only the top eVariant per eGene was used. Only those eQTLs whose 95% confidence interval of the effect size estimate did not overlap zero were used for downstream analysis. To control for differences in power due to eVariant allele frequency, the effect of MAF on eQTL effect size was estimated using LOWESS regression (Matlab function `malowess: span=0.2, robust=true`), and was subtracted from the effect sizes on a per tissue basis.

#### ASE effect size

For each sample, haplotypic expression at all eGenes was calculated by summing counts from all phased, heterozygous SNPs. For a given cis-eQTL variant, assume *xi* is the number of RNA-seq reads aligned to one haplotype, and *y*_*i*_ is the total number of reads aligned to either haplotype in the ith individual. Regulatory effect size of the cis-eQTL was calculated as median log-ratio: *s*(*x,y*) = *median*[log_2_(*x*_*i*_) log_2_(*y*_*i*_ – *x*_*i*_)]. Effect sizes were calculated for cis-eQTLs for which 10 or more individuals with *y*_*i*_ ≥ 10, and the effect sizes were constrained to be less than 100 fold (|s(*x*,*y*)| < log_2_ 100). Confidence intervals for the effect sizes were derived using BCa bootstrap with 100 samples.

### cis-eQTL fine-mapping

#### CaVEMaN

We utilized CaVEMaN (Causal Variant Evidence Mapping with Non-parametric resampling) to estimate the probability that an eVariant was a causal variant (Brown et al., in preparation). We used a non-GTEx reference cis-eQTL dataset from subcutaneous adipose tissue, lymphoblastoid cell lines, skin and whole blood, to simulate causal variants with characteristics matching genuine cis-eQTLs^61^ (effect size, residual variance, minor allele frequency, and distance to the TSS). For each simulation, we calculated the proportion of times the simulated causal variant was among the ith most significant eVariants and denoted this proportion as *p*_*i*_. For each lead eVariant in GTEx, we generated a single-signal expression phenotype by controlling for all covariates fitted in the cis-eQTL mapping and all other eVariants for the gene except the eVariant whose signal we wished to preserve. These data were sampled with replacement 10,000 times and cis-eQTL mapping was performed on each resample. The proportion of times a given eVariant was ranked *i* was calculated, denoted *F*_*i*_. The CaVEMaN score is then defined as 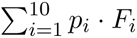. To calibrate CaVEMaN scores, across all genes and tissues simulated (removing blood as an outlier) we divided the CaVEMaN scores of the peak variants into twenty quantiles. Within each quantile, we calculated the proportion of times the lead variant was the causal variant and then drew a monotonically increasing smooth spline from the origin, through the 20 quantiles, to the point (1, 1) using the gsl interpolate functions with the steffen method (gsl-2.1, https://www.gnu.org/software/gsl/). This function provides our mapping of CaVEMaN score of the lead SNP onto the probability it is the causal variant, calibrated using the simulations.

#### CAVIAR

CAVIAR (CAusal Variants Identification in Associated Regions)^29^ uses LD structure to model the observed marginal test statistics for each eGene as following a multivariate normal distribution (MVN). Applying this model, CAVIAR can define a credible set containing all causal variants with probability **ρ*.* To define these credible sets in each tissue, we used a threshold of **ρ* =* 90%. We utilized eCAVIAR (eQTL and GWAS CAVIAR) to colocalize GWAS and eQTL studies for detection of the target genes and relevant tissues^42^. eCAVIAR computes a posterior probability of a variant identified as causal in both GWAS and eQTL studies. We used a cut-off of 1% for colocalization posterior probability based on observations from previous simulations^42^.

To test for a significant relationship between tissue sample size and the size of the 90% credible set, we compared credible set sizes for the top 100 single-tissue cis-eQTLs across tissues (Extended Data Fig. 11a). We further combined cis-eQTL sharing results from METASOFT with CAVIAR's 90% credible sets to test if tissues with shared cis-eQTLs could be used to fine-map the causal variant. Here, from the initial METASOFT results, we identified the top shared cis-eQTL for each eGene by selecting the cis-eQTL with the smallest RE2 *P*-value. For eGenes that had a shared cis-eQTL or a tissue-specific cis-eQTL, we compared the intersection of the 90% credible sets within and between each group (Extended Data Fig. 11b).

### Overlap of tissue-specific and tissue-shared eGenes with disease genes

For each gene tested for multi-tissue eQTLs using METASOFT, we calculated the proportion of tissues for which the gene had a strong eQTL effect (i.e. the proportion of tissues with m-value ≥ 0.9). We defined tissue-specific eGenes as genes in the bottom 10% of the empirical distribution of this proportion. Similarly, we defined tissue-shared eGenes as genes in the top 10% of this distribution. We examined the enrichment of tissue-specific and tissue-shared eGenes in six different gene lists: the NHGRI-EBI GWAS Catalog^62^, the Online Mendelian Inheritance in Man (OMIM) database^63^, the Orphanet database, the ClinVar database^64^, the list of genes with clinically actionable variants reported by the American College of Medical Genetics (ACMG)^65^, and the list of LoF intolerant genes from ExAC^31^. For the GWAS catalog, we restricted to only genes with reported associations. LoF intolerant genes were defined as those with a pLI score ≥ 0.9 in ExAC^31^. We calculated odds ratios and 95% confidence intervals using Fisher’s exact test for both tissue-specific and tissue-shared eGenes in each gene list. For the tissue-specific eGenes, we used as a background the remaining set of genes tested in METASOFT that were not classified as tissue-specific eGenes. Similarly, for tissue-shared eGenes, we used as a background the set of genes not classified as tissue-shared eGenes.

### GWAS analysis

We have previously described the Regulatory Trait Concordance (RTC) score to assess whether a GWAS variant is tagging the same functional variant as a regulatory variant^34^. Briefly, for a cis-eQTL and GWAS variant located in the same region between recombination hotspots, we correct the eQTL phenotype (i.e., gene expression) for all the *N* variants within the region using linear regression, creating *N* pseudo-phenotypes from the residuals of the linear regression. We then test for eQTL association between the cis-eQTL variant and the N pseudo-phenotypes. These *P*-values are subsequently sorted (descending) and ranked, and the rank of the *P*-value arising from the cis-eQTL and GWAS variant corrected phenotype association is found and the score is defined as (*N −* GWASrank) / *N*. The RTC score ranges from 0 to 1 with 1 indicating higher likelihood of shared functional effect.

### CAD GWAS

Data on coronary artery disease and myocardial infarction have been contributed by CARDIo-GRAMplusC4D investigators and have been downloaded from www.cardiogramplusc4d.org.

## Data availability

Genotype data from the GTEx V6p release are available in dbGaP (study accession phs000424.v6.p1; www.ncbi.nlm.nih.gov/projects/gap/cgi-bin/study.cgi?study_id=phs000424.v6.p1). The VCFs for the imputed array data are in phg000520.v2.GTEx MidPoint Imputation.genotype-calls-vcf.c1.GRU.tar (the archive contains a VCF for chromosomes 1-22 and a VCF for chromosome X). Allelic expression data is also available in dbGap. Expression data (read counts and RPKM) and eQTL input files (normalized expression data and covariates for 44 the tissues) from the GTEx V6p release are available from the GTEx Portal (http://gtexportal.org). eQTL results are available from the GTEx Portal. In addition to results tables for the 44 tissues in this study (eGenes, significant variant-gene pairs, and all variant-gene pairs tested), the portal provides multiple interactive visualization and data exploration features for eQTLs, including:

- eQTL box plot: displays variant-gene associations
- Gene eQTL Visualizer: displays all significant associations for a gene across tissues and linkage disequilibrium information
- Multi-tissue eQTL plot: displays multi-tissue posterior probabilities from meta-analysis against single-tissue association results
- IGV browser: displays eQTL across tissues and GWAS Catalog results for a selected genomic region

## Acknowledgments

The Genotype-Tissue Expression (GTEx) project was supported by the Common Fund of the Office of the Director of the National Institutes of Health (commonfund.nih.gov/GTEx). Additional funds were provided by the National Cancer Institute (NCI), National Human Genome Research Institute (NHGRI), National Heart, Lung, and Blood Institute (NHLBI), National Institute on Drug Abuse (NIDA), National Institute of Mental Health (NIMH), and National Institute of Neurological Disorders and Stroke (NINDS). Donors were enrolled at Biospecimen Source Sites funded by NCISAIC-Frederick, Inc. (SAIC-F) subcontracts to the National Disease Research Interchange (10XS170) and Roswell Park Cancer Institute (10XS171). The Laboratory, Data Analysis, and Coordinating Center (LDACC) was funded through a contract (HHSN268201000029C) to The Broad Institute, Inc. Biorepository operations were funded through an SAIC-F subcontract to Van Andel Institute (10ST1035). Additional data repository and project management were provided by SAIC-F (HHSN261200800001E). The Brain Bank was supported by a supplement to University of Miami grant DA006227.

We thank Anshul Kundaje and Oana Ursu for input on the HI-C analysis. J.R.D. is supported by a Lucille P. Markey Biomedical Research Stanford Graduate Fellowship. J.R.D., Z.Z., and N.A.T. acknowledge the Stanford Genome Training Program (SGTP; NIH/NHGRI T32HG000044). Z.Z. is also supported by the National Science Foundation (NSF) GRFP (DGE-114747). L.F. is supported by the Stanford Center for Computational, Evolutionary, and Human Genomics (CEHG). D.G.M. is supported by a “la Caixa”-Severo Ochoa pre-doctoral fellowship. D.M. is supported by NIH grants U54DK105566 and R01GM104371. B.J.S is supported by NIH training grant T32 GM007057. E.K.T is supported by a Hewlett-Packard Stanford Graduate Fellowship and a doctoral scholarship from the Natural Science and Engineering Council of Canada. T.S. is supported by a National Science Foundation Graduate Research Fellowship (DGE-1656518). A.B. is supported by the Searle Scholars Program and NIH grant 1R01MH109905. A.B., C.D.Bu. and S.B.M are supported by NIH grant R01HG008150 (NHGRI; Non-Coding Variants Program). S.B.M. and C.D.Bu. are supported by NHGRI grants U01HG007436 and U01HG009080. A.B., C.D.Bu., E.T.D., S.E.C., T.L, and S.B.M. are supported by NIH grants R01MH101814 (NIH Common Fund; GTEx Program). C.D.Br., Y.P., and B.E.E. are supported by NIH grant R01MH101822. G.L., A.B.N, J.J.P., A.S., Y-H.Z., and F.A.W. are supported by NIH grants R01MH101819, R01HG009125, and R21HG007840. T.L. and P.M. are supported by NIH grant R01MH106842. T.L. is supported by the NIH grant UM1HG008901. T.L. and S.E.C. are supported by the NIH contract HHSN2682010000029C. C.S. and C.B.P. are supported by NIH grant R01 MH101782.

## Contributions

F.A., A.V.S., N.J.C., G.G., K.G.A., and E.T.D. contributed to study design. F.A., A.A.B., S.E.C., J.R.D., P.M., A.V.S., Z.Z., N.S.A., L.F., E.R.G., E.G., M.J.G., Y.H., F.H., X.L., X.Li., B.L., D.G-M., H.O., J.J.P., Y.P., C.B.P., G.Q., S.R., A.A.S., T.C.S., B.J.S., T.J.S., N.A.T., E.K.T., H.Z., Y-H.Z., A.B., C.D.Bu., B.E.E., E.E., M.K., G.L., D.G.M., A.B.N., C.S., X.W., F.A.W., T.L., K.G.A., E.T.D., C.D.Br., and S.B.M. contributed analysis. C.D.Br. and S.B.M. wrote the manuscript. F.A., A.A.B., S.E.C., J.R.D., P.M., A.V.S, Z.Z., C.B.P., B.J.S., A.B., C.S., T.L., K.G.A., E.T.D., C.D.Br., and S.B.M. contributed text to the manuscript. All authors reviewed and revised the manuscript.

## Corresponding authors

Emmanouil T. Dermitzakis, Christopher D. Brown, Stephen B. Montgomery

